# Investing in Open Science: Key Considerations for Funders

**DOI:** 10.1101/2024.12.09.627554

**Authors:** Dana E. Cobb-Lewis, Devin Synder, Sonya Dumanis, Robert Thibault, Barbara Marebwa, Elisia Clark, Lara St. Clair, Leslie Kirsch, Michelle Durborow, Ekemini Riley

## Abstract

The open science movement aims to transform the research landscape by promoting research transparency in order to enable reproducibility and replicability, lower the barriers for collaboration, and reduce unnecessary duplication. Recently, in recognition of the value of open science, funding agencies have begun to mandate open science policies as a condition in grantee awards. However, operationalization and implementation of an open science policy can have unanticipated costs and logistical barriers, which can impact both the funder, as well as the grantee. These factors should be considered when implementing an open science policy.

The Aligning Science Across Parkinson’s (ASAP) initiative utilizes a comprehensive open science policy, which, in addition to requiring immediate free online access to all publications, also requires all newly-generated datasets, protocols, code, and key lab materials be shared by the time of publication. Moreover, preprints must be posted to a preprint repository by the time of manuscript submission to a journal for review.

Here, we outline the potential costs associated with implementing and enforcing this open science policy. We recommend that funders take these considerations into account when investing in open science policies within the biomedical research ecosystem.

## Introduction

Open science is a multifaceted movement that aims to make the processes and outputs of scientific research findable, accessible, and reusable, with the goal of accelerating the pace of scientific discovery by allowing others to build upon the work of those who came before. Open science is transforming the research landscape, with funders and journals mandating varying levels of openness ^1–5^. However, developing operational workflows to promote the adoption and implementation of open science policies can pose unanticipated costs and logistical barriers to funders. The Aligning Science Across Parkinson’s (ASAP) initiative is a coordinated research program designed to accelerate discoveries for Parkinson’s disease through facilitating collaboration, generating research enabling resources, and data sharing. Since its inception, the initiative has embraced a comprehensive [open science policy](https://parkinsonsroadmap.org/open-science-policy/#) with five requirements ^6,7^:

**Requirement 1:** Share all research outputs by time of publication.
**Requirement 2:** Identify all research inputs in final publication.
**Requirement 3:** Ensure immediate open access.
**Requirement 4:** Acknowledge ASAP.
**Requirement 5:** Share outputs with the ASAP network.

Here, we examine the feasibility and sustainability of adopting this open science policy from a funder’s perspective. We break down each of the five overarching requirements in the [ASAP Open Science Policy](https://parkinsonsroadmap.org/open-science-policy/#), outlining the barriers and costs associated with implementing and enforcing the policy (Table 1).

**Table 1.**
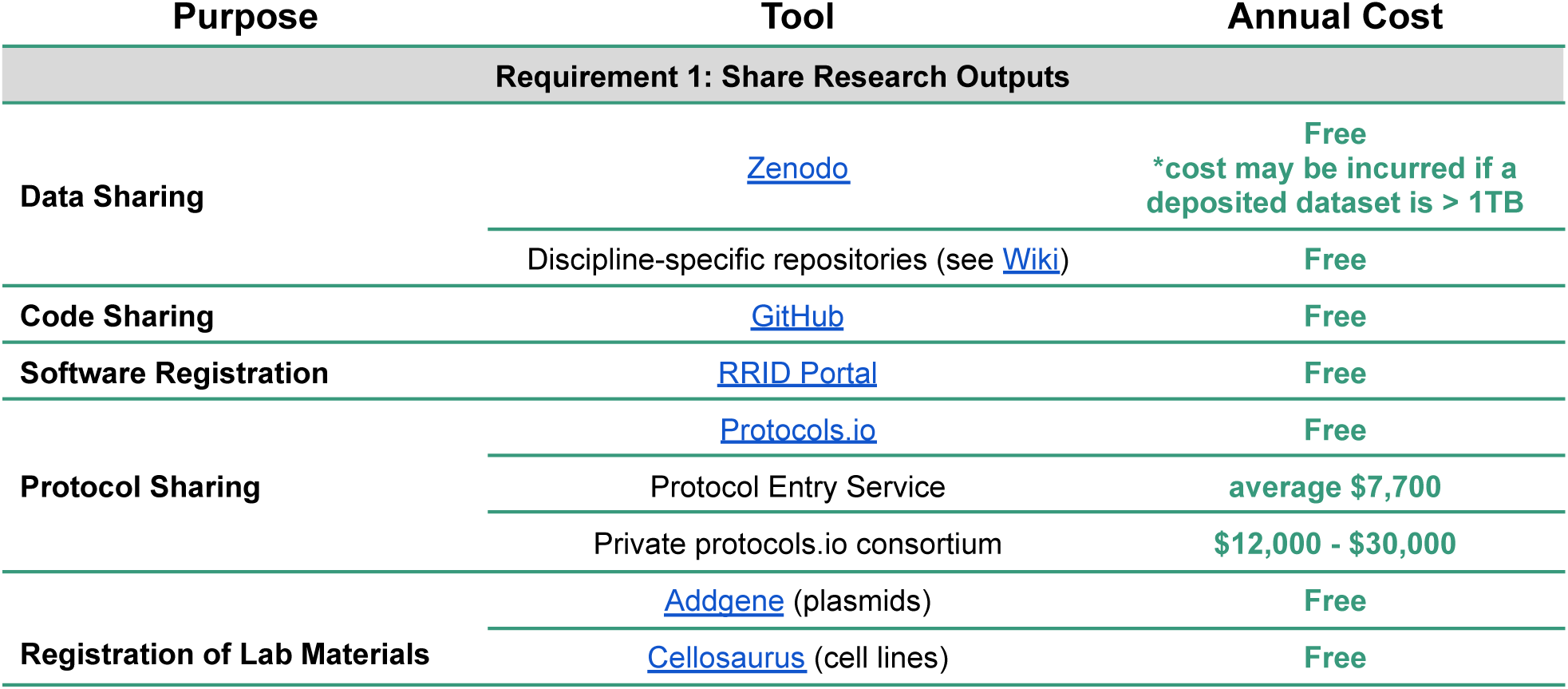
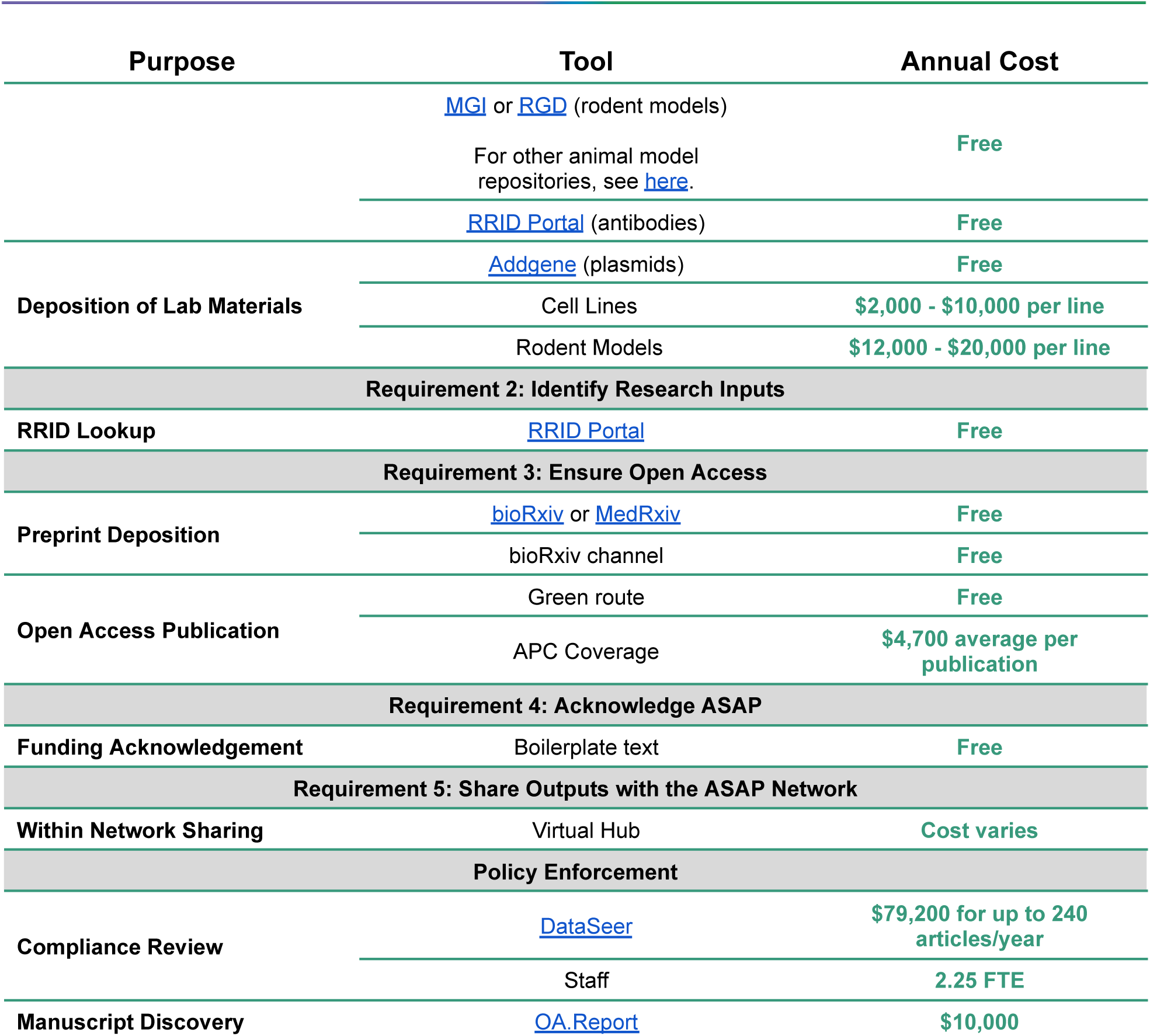
Costs associated with implementing an open science policy. This table outlines the estimated annual cost of implementing and enforcing each aspect of the ASAP Open Science Policy using repositories and resources recommended by ASAP. The table is broken down by each requirement of the policy. Note that the largest expenditure comes from supporting lab material deposition, covering article processing charges, and policy enforcement. Costs associated with lab material deposition vary based on CROs and contract negotiations. All amounts are listed in USD.

## Requirement 1: Share Research Outputs

ASAP defines **research outputs** as any data, code & software, protocols, or key lab materials that are generated during a research study ^6,7^. Several funding agencies are moving towards mandating that data be shared as part of their open science policy ^8–10^. The [ASAP Open Science Policy](https://parkinsonsroadmap.org/open-science-policy/#) goes beyond a data sharing mandate, requiring all research outputs (data, code & software, and protocols) generated as part of an ASAP-funded study to be deposited in a community-recognized repository by the time of publication, with a license that allows for reuse. Any accompanying information (e.g., README files) necessary to facilitate reuse of research outputs must be included in the deposit and the associated persistent identifier must be findable via the publication. In the early years of the ASAP Initiative, we required deposition of all newly-generated lab materials by time of publication. However, our grantees were rarely able to meet this requirement due to the lengthy process associated with depositing lab materials. In response to grantee feedback, we have modified our policy to require *registration* and process of deposition initiated rather than deposition of the lab material output by the time of publication.

### Data

#### Barriers to implementation

From a funder’s perspective, there are several implementation barriers when enforcing a data sharing policy. Despite growing mandates for data sharing, datasets are often not shared in publicly accessible repositories, with one meta-analysis finding only 2% of medical and health sciences publications had associated publicly available data ^11^. Some researchers believe that sharing data “upon request” or as supplemental files is sufficient. However, numerous studies have shown that, despite “data available upon request” statements in their papers, researchers are rarely willing or able to share their data upon request ^12–18^. Additionally, supplemental files are often subject to link rot and content drift, underlining the importance of public repositories meeting best practices for the long-term preservation of data ^19–22^. Perhaps not surprisingly, data from publications in journals with mandated data sharing policies are significantly more likely to be findable online compared to data from publications in journals with no mandate ^23^. While the easiest mechanism for changing this cultural norm may rely on clear data sharing mandates from journals themselves, this type of requirement is not widespread. Therefore, for now, grantee training and education about the importance of sharing data in public repositories for long-term preservation of data may help generate buy-in for a data sharing policy. The ASAP initiative has committed to providing resources, such as Open Science Guides and a [Data Repository Wiki](https://youneedawiki.com/app/page/1dTXSFCM-EmnEMTQjWsYWt6V5hPiAokXt?p=1dTXSFCM-EmnEMTQjWsYWt6V5hPiAokXt), for ASAP grantees, as well as the scientific community at large. However, the creation and maintenance of these resources costs time and effort for the ASAP Open Science Team.

Data curation and data standards are also important considerations. It is not enough for the data to be made available; for the data to be useful, its distribution must include additional support, such as README files and data dictionaries. Ensuring that datasets are well-curated is essential for their meaningful use and interpretation. However, curating datasets requires time, effort, and expertise. Data sharing standards vary across disciplines (or do not exist at all), making it challenging for researchers to navigate where and how to share their data effectively. While the burden of curating data often falls on the individual grantees, we have found that providing educational guidance (including templates and links to metadata standards for data types) helps grantees comply with this policy. Sharing cleaned, tabular data, as compared to non-tabular or raw data, can be easier for grantees. However, sharing tabular data effectively still requires a high degree of data management and documentation, and ultimately, sharing only cleaned data limits the extent to which the data could be aggregated for meta-analysis or reused in future studies.

Addressing these barriers requires concerted effort from stakeholders across the research community, including funders, institutions, publishers, and researchers themselves, to develop standardized guidelines and foster a culture of data sharing and collaboration.

#### Costs associated with data sharing

Data sharing costs for funders fall into three major categories: effective curation, data storage/hosting, and data infrastructure. Effective curation to prepare data for reuse may be performed by either a grantee or a designated data curator; in either case, there is a cost in personnel time and potentially in computational resources. The cost generally depends on the complexity and size of the dataset. Curation costs may be reduced by using a repository which includes curation services, such as [Dryad](https://datadryad.org/stash) or an institutional repository, but these repositories may charge a service fee and may not cover all datatypes. The majority of discipline-specific and generalist data repositories are free to use for storage/hosting and distribution by default (Table 1), however, costs can escalate for larger datasets and specialized platforms. For example, [Zenodo](https://zenodo.org/), the generalist repository utilized by ASAP, provides free storage and distribution up to 1TB, but beyond that, users incur charges. For very large datasets, egress costs for distribution can be significant. Upgrades to, and maintenance of platforms or services may require further financial investment, posing a barrier to researchers, especially those with limited funding. This is a particularly costly challenge for funders and researchers working with human subject data, which may have a higher bar to meet ethical and legal requirements, and often incur additional data sharing costs at every stage of sharing.

Data sharing at scale relies on high quality data and infrastructure ^24^. We have found that it may be beneficial for funders to support and/or develop data infrastructure for the specific data types emerging from their programs. To that end, ASAP partners with multiple organizations to support the AMP-PD data repository, a public-private partnership for Parkinson’s disease datasets. AMP-PD provides a platform to centralize, harmonize, and securely distribute genetic and multi-’omic data relevant to Parkinson’s alongside a standardized minimum set of key clinical data. ASAP also developed a data sharing tool, the [CRN Cloud](https://cloud.parkinsonsroadmap.org/collections), which currently consists of harmonized collections of human postmortem-derived brain sequencing data, a precious resource for the scientific community. The CRN Cloud provides a data platform that allows these data to be stored securely within a cloud environment while still being accessible to the broader research community, and provides a collaborative environment where scientists can access, analyze, and share these datasets. Developing the CRN Cloud has allowed us to more cost-effectively and quickly curate, store, and distribute very large multimodal datasets, with additional collections in curation and scheduled for future distribution based on our strategic priorities. It has also provided an opportunity to enforce data standards, support consistent curation and data quality to enable interoperability between datasets, and develop more effective guidance for data reusers. Note that creating cloud-based resources like the CRN Cloud requires capital, and depending on the features required, can easily become a greater than million dollar line item in a funder’s budget.

Additionally, some disciplines are working towards standardized, non-proprietary file types, but this requires concerted efforts across the field and significant infrastructure support. For example, sharing neurophysiology data remains a challenge due to large file sizes and proprietary file types. Additionally, sharing of this data may require synchronization of neurophysiology data with other data streams (e.g., behavioral outputs). Therefore, we support a collaboration with [Catalyst Neuro](https://www.catalystneuro.com/) to build bespoke conversion and transfer pipelines for neurophysiology data to allow for easy archiving of these datasets onto [DANDI](https://dandiarchive.org/), the NIH BRAIN Initiative-sponsored neurophysiology repository. These curation efforts around building these workflow pipelines amount to around a $30,000 investment per lab we support. Supporting these data infrastructure projects represents a non-trivial cost and a strategic investment towards the preservation, accessibility, and reusability of ASAP-funded data.

Implementation of a data sharing policy may cost the funder nothing. However, this is dependent upon the design of the policy and the commitment on the part of the funder to support data infrastructure that may be needed to house data generated by grantees.

### Code and Software

#### Barriers to implementation

One of the primary barriers to implementing a code sharing policy is inertia. Computational reproducibility is increasingly viewed as a minimum standard for scientific research to meet. Despite this, sharing of analytical scripts is not common practice ^25,26^. Estimates of analytical script sharing in scientific publications vary widely (<0.5% to 34%) and, even when shared, scripts are often insufficiently annotated or incomplete ^11,27,28^. Many researchers may not have received training in best practices for code documentation as part of their technical training, and may not feel they need to improve this skill.

Enforcement of this policy is another barrier to implementation. In our current workflow, the ASAP Open Science Team checks whether or not the analytical scripts are shared, not if they are reusable. Checking that scripts are comprehensive and well-documented takes substantial time and expertise, and may not be feasible for a funder since it often requires access to specific computational resources. Additionally, for many manuscripts where analytical scripts are *not* generated due to the technology used (e.g., Excel, GUI-based analytics tools), specific details of cleaning, analysis, and visualization of data often goes unreported. To address this, we recently began recommending that grantees write instructions for actions they perform in point-and-click software; however, standards, best practices, and examples need to be established in order to codify this as mandatory policy.

Addressing these barriers requires a concerted effort from stakeholders across the research community to develop standardized guidelines, and foster a culture of sharing all analytical scripts and assigning all code deposits a persistent identifier.

#### Costs associated with sharing code and software

Requirements to share analytical scripts in a publicly accessible repository do not generally have any associated funder costs. Although paid plans offer more storage and technical support, which grantees may wish to take advantage of, it is free for grantees to share their code in a generalist repository such as [GitHub](https://github.com/). Additionally, licensing scripts is free and easy to achieve by generating a LICENSE file for the GitHub repository.

Sharing of open-source executable software or software tool libraries/packages is more complicated. Like analytical scripts, licensing of open-source software is free and source code can be distributed via Github or language libraries. Additionally, registration of software via [RRID Portal](https://rrid.site/) is also free. However, maintaining software comes with associated costs, especially if software needs to be debugged and/or upgraded due to high popularity/usage or if data storage is required. This cost often falls to the grantee, but in some cases it may be a worthwhile strategic investment for funders to offset these costs if the software tool is of particular interest or use.

### Protocols

#### Barrier to implementation

Funders that require grantees to share detailed protocols through a public repository should recognize that there are several barriers to implementing this policy. One of the main barriers is that grantees do not share sufficiently detailed protocols with researchers outside of their lab. In a small survey of researchers across disciplines, nearly a quarter (23%) of respondents reported not sharing protocols or detailed methods outside of their lab or group and only a third (33%) indicated that they share their protocols in a written format ^29^.

Another consideration for funders implementing a protocol sharing requirement is the time it takes for grantees to create detailed protocols and to learn how to use a protocol repository. In a survey of researchers, the majority (74%) reported not sharing protocols due to not knowing where to share detailed methods and the effort and time it takes to create detailed methods ^29^. In an effort to assist grantees, ASAP has provided step-by-step instructions for how to use the protocol repository protocols.io and how to write a recipe-style protocol. Additionally, we cover fees associated with using the protocols.io [Protocol Entry Service](https://www.protocols.io/help/protocol-entry-service), which allows grantees to submit their protocols to be formatted and imported by the protocols.io team.

Addressing these barriers requires education and may involve strategic investment from funders to support development and sharing of recipe-style protocols.

#### Costs associated with protocols.io

In general, the costs that a funder might encounter for implementing a protocol sharing requirement can be relatively minimal. However, costs are highly dependent upon how much support the funder provides to the grantee. Some funders may choose to utilize the free version of protocols.io, which gives users the ability to create an unlimited amount of public protocols (for example, see the [CZI Neurodegeneration Challenge Network](https://www.protocols.io/workspaces/neurodegeneration-method-development-community1) community, with over 100 protocols shared).

ASAP grantees wanted the ability to share private protocols internally with the network before making them public, as there were concerns about publishing a protocol prior to a grantee completing troubleshooting. In light of this concern, we opted to support the creation of a paid, private ASAP workspace in protocols.io for members of the Collaborative Research Network (CRN). Nearly 500 members of the ASAP workspace actively utilize this platform, which houses over 2000 protocols, to create private protocols that are shared internally with the ASAP CRN prior to publication. Note that a majority of these private protocols do become public, with over 1400 (70%) of protocols currently public. Maintaining a private workspace for grantees through protocols.io can be costly, ranging from $12,000-$30,000 per year depending on the tier and support provided to grantees. In addition, we cover fees associated with using the protocols.io [Protocol Entry Service](https://www.protocols.io/help/protocol-entry-service). Our grantees utilize the Protocol Entry Service an average of 152 times per year. Each entry costs $50, representing an average annual cost of $7,600 (Table 2).

**Table 2.**
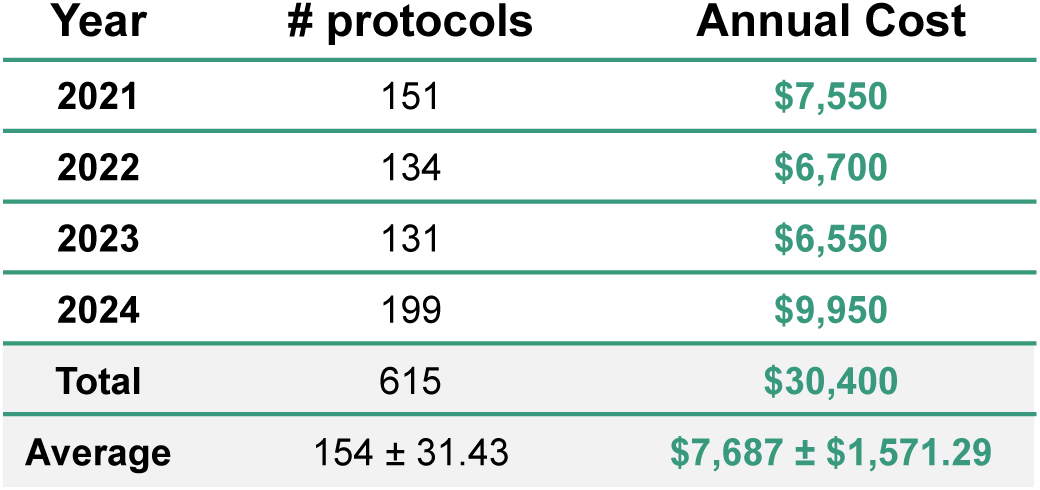
Costs associated with Protocol Entry Service. The Protocol Entry Service formats, edits, and publishes protocols to protocols.io on behalf of grantees. Each entry is $50. All amounts are listed in USD and averages are reported as mean ± SD.

| Year | # protocols | Annual Cost |
| --- | --- | --- |
| 2021 | 151 | \$7,550 |
| 2022 | 134 | \$6,700 |
| 2023 | 131 | \$6,550 |
| 2024 | 199 | \$9,950 |
| **Total** | 615 | **\$30,400** |
| **Average** | 154 ± 31.43 | **\$7,687 ± \$1,571.29** |

### Lab Materials

#### Barriers to implementation

There are a couple considerations for funders to be mindful of when requiring grantees to register and/or deposit lab materials. The time it takes to deposit lab materials at a repository is a major barrier to implementing this policy, particularly when it comes to deposition by time of publication. The time to deposit a new lab material ranges from 6 – 24 months to finalize agreements, import material, perform quality control, and prepare for distribution (Table 3). There are often delays related to communicating with the Tech Transfer Office of the depositing investigator’s institution or missing information. Therefore, in order to ensure that a lab material is registered and deposited by time of publication, investigators should start the deposition process at the pre-publication stage.

**Table 3.** Activities and timelines associated with lab material deposition. The MJFF Research Tools Program assists grantees with deposition of newly-generated lab materials, including rodent models, iPSCs, and plasmids. The Tools Program does not currently support deposition of antibodies. The timeline column encompasses the “covered activities,” however, time to distribution may take up to 24 months to finalize legal agreements.

| Tool | Information to Provide | Material to Ship | Covered Activities | Timeline |
| --- | --- | --- | --- | --- |
| Rodent | • Mutation(s) and transgene(s)<br>• Genetic background<br>• Phenotype<br>• Genotyping assay<br>• Associated publications | 1 mouse | • Shipping and transport<br>• Line rederivation<br>• Cryopreservation<br>• Colony generation<br>• Product page<br>• Maintenance as a live colony for 1 year | 6-9 months |
| Induced pluripotent stem cells (iPSC) | • Mutation(s)<br>• Genetic background<br>• Phenotype<br>• Quality control<br>• Associated publications | 12 vials of cells | • Shipping<br>• Storage<br>• Line expansion<br>• Cryopreservation<br>• Product page<br>• Maintenance | 3-6 months |
| Plasmids | • Vector maps and components | 15 uL DNA or bacterial streaks | • Prepaid shipping envelope, small boxes, and barcode labels | 1-3 months |

There are also several legal and ethical considerations when implementing a policy requiring lab materials to be deposited and/or registered. A detailed history of source material (i.e. background, original creator, mutations, transfer agreements, and previous crosses) used to generate lab materials are required to protect the intellectual property (IP) rights of the investigator and institution of material origin. This is often overlooked and can delay or prohibit deposition or registration of lab materials for sharing with the wider research community. Other considerations include confirming proper consent for generation and sharing of human-derived cell lines, Material Transfer Agreements (MTA) that allow deposition and registration of materials by institutions beyond the original one, and licensing barriers for non-profit and for-profit organizations. We have found that having a Tools Program dedicated to walking grantees through this process is essential for funders who would require that lab materials be made commercially available as part of their open science policy.

Addressing these barriers requires education and may involve strategic investment from funders to support deposition and registration of lab materials.

#### Costs associated with registration and deposition of lab materials

Registration of newly-generated lab materials can be done at no cost through resource information networks and databases, such as [RRID Portal](https://rrid.site/), [Cellosaurus](https://www.cellosaurus.org/), and [MGI](https://www.informatics.jax.org/mgihome/submissions/amsp_submission.cgi) (see Table 1). Deposition is ideal compared to registration since it also allows the Contract Research Organization (CRO) to manage orders and maintain quality for distribution. However, supporting deposition of lab materials can be very costly, ranging from $0 - $20K per tool (Table 1). ASAP has partnered with the [MJFF Research Tools Program](https://www.michaeljfox.org/sponsored-tools-program) to help facilitate and oversee this process. Although running a Tools Program represents a further financial investment in addition to the costs associated with deposition itself, grantees benefit from utilizing the Tools Program to provide support and guidance during the deposition process.

## Requirement 2: Identify Research Inputs

ASAP defines **research inputs** as any data, protocols, code & software, or key lab materials that are *not* generated in a manuscript (i.e., inputs are re-used outputs from previous work) ^6,7^. Unambiguously identifying research inputs, particularly protocols, software, and lab materials, has been shown to be critical for replicability and reproducibility of science ^30–33^. Journals have taken note of the need for transparency when it comes to sharing of lab materials, with several (e.g., eLife and Cell Press) requiring the inclusion of Key Resource Tables in publications, which detail lab materials used in a study and their associated persistent identifiers ^34,35^.

The [ASAP Open Science Policy](https://parkinsonsroadmap.org/open-science-policy/#) requires that all data, software, protocols, and key lab materials used in a study–but which were not generated as part of an ASAP-funded study–are unambiguously identified in the study’s publication.

### Barriers associated with implementation

From the funder’s perspective, there are few barriers to implementation of an open science policy requiring identification of research inputs with persistent identifiers. The primary barrier to implementation is a lack of education from grantees about what should be shared. For example, many grantees share catalog numbers, however this is not sufficient to uniquely identify the material, as vendors may merge and catalog numbers can change ^31^. We have found that requiring Key Resource Tables (KRT; see the ASAP Open Science Guides for a template) in a manuscript is a useful tool to facilitate sharing of research inputs and specifying the precise type of information grantees are being asked to share.

### Costs associated with identifying research inputs

Requirements to share research inputs do not have any associated funder costs. Grantees can use platforms like [RRID Portal](https://rrid.site/) for free to search for persistent identifiers associated with plasmids, cell lines, antibodies, organisms, and software tools.

## Requirement 3: Ensure Open Access

Following the Budapest Open Access Initiative (BOAI) declaration in the early 2000s, open access mandates have been implemented by funders in many countries ^4,36–39^. Although open access is not yet the default publishing model, the number of scientific publications available via open access has grown since open access mandates first began, and some subscription journals are transitioning to open access models ^40–42^. Researchers are also taking advantage of preprints, with submissions to the preprint server bioRxiv steadily increasing since its inception in 2013 ^43,44^.

The [ASAP Open Science Policy](https://parkinsonsroadmap.org/open-science-policy/#) requires preprints to be posted to a preprint repository no later than the date a manuscript is submitted to a journal for review. Publications must be immediately publicly available with no embargo. Additionally, both preprints and publications must be licensed for reuse with a CC BY 4.0 or CC0 license.

### Preprints

#### Barriers to implementation

From a funder’s perspective, the primary barrier to implementing a preprint policy is addressing concerns from grantees about spreading misinformation or low quality science, and potential for authors to be “scooped” ^45–47^. Despite these concerns, the majority of papers posted to the preprint servers [bioRxiv](https://www.biorxiv.org/) and [MedRxiv](https://www.medrxiv.org/) are subsequently published in a peer reviewed journal with no significant differences in the content and experimental results between the preprint and publication ^43,48–50^. Additionally, the majority of preprint authors (99.3%) report no problems with scooping ^48,51–53^. Preprints provide date-stamped priority claims and establish intellectual precedence, with some journals even offering scoop protection policies that honor priority claims of preprints ^53–55^.

Addressing this barrier requires concerted effort from stakeholders across the research community, including funders, to foster a culture of sharing research early and often.

#### Costs associated with preprint deposition

Requirements to post preprints to a preprint server do not have any associated funder costs. Posting a preprint is free via a number of preprint servers such as [arXiv](https://arxiv.org/), [bioRxiv](https://www.biorxiv.org/), and [medRxiv](https://www.medrxiv.org/). Funders can create a [channel](https://connect.biorxiv.org/relate/content/195), a curated collection of preprints from bioRxiv and medRxiv that share the subject area of a particular organization (e.g., [Aligning Science Across Parkinson’s](https://connect.biorxiv.org/relate/content/195)) on bioRxiv for no cost. However, the funder is responsible for curation of these channels, by either manually entering each preprint associated with their channel or by submitting a list of preprint DOIs, which represents a small associated labor cost.

### Publications

#### Barriers to implementation

From a funder’s perspective, there are two ways to approach an open access publication requirement: (1) funders may choose to provide coverage of article processing charges (APCs) for publishing in gold or hybrid open access journals, or (2) funders may choose to promote the “green route” whereby the “author accepted manuscript” is deposited to a community-accepted repository (e.g., self-archiving). Should a funder elect to pay for APCs, they should be aware that this route requires funders to establish a standardized workflow for processing payments. Using automation and/or collaborating with publishers for invoice payment can reduce administrative burden. Additionally, education of grantees about publishing open access, especially with regard to open access licenses is needed. We have found that many journals treat review articles differently than original research articles, such that a journal that appears to be open access through the [Sherpa Romeo tool](https://www.sherpa.ac.uk/romeo/) may not be for review articles. This results in review articles being hidden behind a paywall, and requires effort on the part of the author to share the review via the green route after an embargo period. Education and clear guidelines for grantees on how to approach review articles is needed.

#### Costs associated with ensuring open access

If funders choose to pay APCs to ensure publications are open access via gold or hybrid journals, costs can vary widely, with higher impact journals typically charging the highest APCs (Figure 1). These costs can be offset if organizations or institutions have memberships with publishers to provide authors with discounted or waived APCs. Additionally, some open access journals offer fee waivers or discounts for authors from low-income countries, students, early-career researchers, or others who may have difficulty covering full publication costs.

**Figure 1.**
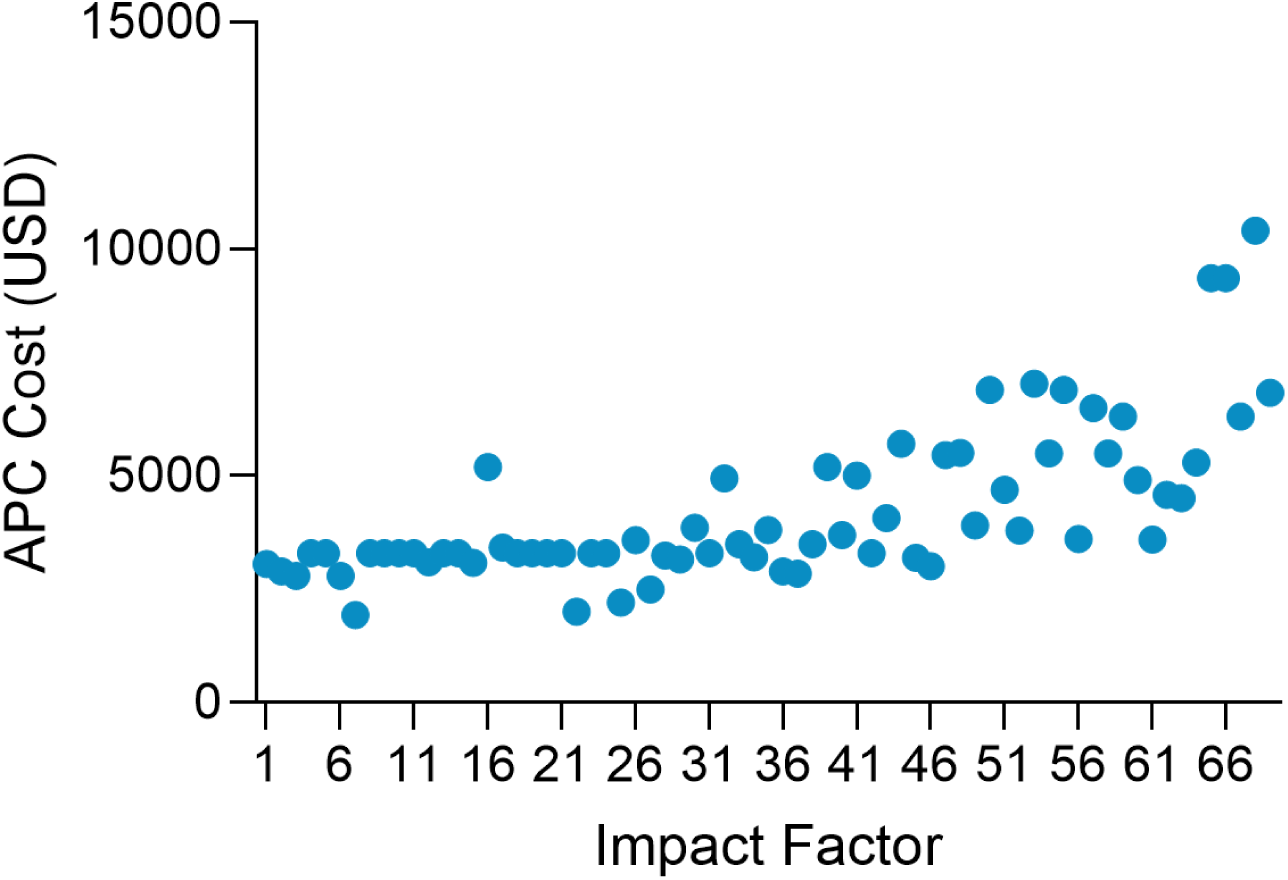
Association of journal impact factor and article processing charge (APC) costs. APCs and impact factors for the most common journals for ASAP publications. Journals with the highest impact factor have the highest APCs. All amounts are listed in USD.

In order to facilitate open access publishing, we offer APC coverage for ASAP-funded publications in addition to the grantee research budget. The number of publications for which we have covered APCs has grown since 2021 (Table 4). On average, we spend $4,700 per publication, investing over $669,000 since 2021 (Table 4).

**Table 4.** Article processing charge expenditures associated with the ASAP Open Science Policy. APCs paid for ASAP-funded publications were tracked over time. All amounts are listed in USD and averages are reported as mean ± SD.

| Year | # Publications Covered | Average APC Cost | Total APC Cost |
| --- | --- | --- | --- |
| 2021 | 7 | \$3,496.14 ± \$1,253.83 | \$24,473.00 |
| 2022 | 40 | \$4,120.98 ± \$1,564.97 | \$164,839.00 |
| 2023 | 40 | \$5,399.36 ± \$3,455.16 | \$215,974.38 |
| 2024 | 54 | \$ 4,901.41 ± \$2,491.93 | \$264,676.64 |
| All Years | 141 | \$4,751.51 ± \$2,597.51 | \$669,963.02 |

Funding APC coverage is a strategic investment we have undertaken in our commitment towards advancing open science and to increase adoption of our policy by grantees. We have paid APCs for about 43% (141/325) of all ASAP-funded publications to date. In the early days of the program, we paid APCs for all ASAP-funded manuscripts to ensure open access. However, in 2023, we began applying stricter eligibility criteria, requiring that manuscripts must be fully compliant with the ASAP Open Science Policy to be eligible for APC coverage. This policy change has further incentivized grantees to adhere to the ASAP Open Science Policy. However, this policy change also requires extensive compliance checks by ASAP staff to ensure the final publication is compliant.

While providing APC coverage offers many benefits, APC costs continue to rise, therefore, offering this coverage involves a substantial financial commitment on the part of the funder ^56–58^. Funders may choose to require self-archiving in place of publishing in open access journals. This method has a financial advantage but funders should know that if a manuscript is published under an embargo period, self-archiving cannot happen until after the embargo is lifted, and some journals do not permit authors to self-archive ^59,60^. However, new policies are changing the conversation surrounding self-archiving ^61^. Publishing in platinum or diamond journals is another APC-free publishing pathway, however only 4.3% of diamond journals are compliant with the coalitionS PlanS Open Science Standards, an initiative committed to full and immediate open access publishing ^62^. This suggests that these journals are not yet an ideal route for publishing open access, but as they grow and receive more support, diamond journals could become a viable option for free open access publishing ^63–65^.

Some funders are approaching open access publishing in yet another way, by choosing to require only preprint deposition as a part of their open science policy, entirely bypassing the costs and politics associated with the publishing world. For example, the Gates Foundation recently made waves when they announced the launch of their own preprint server for Gates Foundation grantees, [VeriXiv](https://verixiv.org/), in collaboration with F1000 ^66^. Beginning in 2025, the Gates Foundation will require preprint deposition to VeriXiv and will no longer pay APCs for publications ^67^.

## Requirement 4: Acknowledge ASAP

Acknowledgement of funding sources enhances the research process by ensuring transparency and mitigating conflicts of interest ^68^. From a funder’s perspective, requiring funding acknowledgements is important for tracking the number of research publication outputs associated with a funder, as well as for assessing impact and tracking collaboration ^68,69^.

The [ASAP Open Science Policy](https://parkinsonsroadmap.org/open-science-policy/#) requires that manuscripts and other research outputs that were partially or fully funded by ASAP must acknowledge ASAP. In addition, all ASAP-affiliated authors must share their ORCID and include ASAP as an affiliation.

### Barriers to implementation

When implementing a funding acknowledgement into an open science policy, funders should recognize that there may be confusion from grantees about what to include in a funding acknowledgement and when to include an acknowledgement ^68^. In order to facilitate compliance with this policy, we provide boilerplate text for grantees to use in their funding acknowledgement and affiliation. We also provide resources for grantees that explain how to determine if their manuscript should be considered ASAP-funded or not, and provide staff support to determine this as well. Although the requirement to acknowledge funding in preprints and publications is relatively easy for grantees to comply with, grantees are less likely to include funding acknowledgements in their research outputs such as a dataset or protocol, etc. Education and training can help improve compliance with this aspect of the policy.

Finally, if requiring funding acknowledgements, funders will need to ensure they have a research organization identifier from the [Research Organization Registry](https://ror.org/) (ROR), as most repositories and publishing platforms use the ROR to provide dropdown options for their users to select the funding agencies to be acknowledged.

### Costs associated with funding acknowledgements

Requirements to include funding acknowledgements do not have any associated funder costs.

## Requirement 5: Share Outputs with the ASAP Network

In an effort to encourage knowledge and resource sharing, ASAP collaborated with [YLD](https://www.yld.io/), a software engineering and design consultancy group, to create the ASAP CRN and GP2 Hubs; private, virtual platforms for grantees. The Hub provides grantees with a research output management system, a space to share their research outputs and findings with the larger ASAP network even earlier than a preprint would allow, further accelerating discovery and increasing collaboration.

The [ASAP Open Science Policy](https://parkinsonsroadmap.org/open-science-policy/#) requires that all ASAP-funded research outputs, including manuscripts, be shared on the ASAP grantee virtual platform, the ASAP Hub, no later than time of publication. While sharing research outputs on the Hub provides an avenue for grantees to learn more about available resources, it also acts as a source for sharing ASAP-funded outputs with the scientific community at large. Any research output on the Hub that is both ASAP-funded and public is also added to our publicly available [ASAP catalog](https://parkinsonsroadmap.org/catalog/#) via an automated API between the two platforms. As of October 2024, the external ASAP catalog showcased over 400 articles, 100 scripts, 200 datasets, 375 lab materials, and 875 protocols.

### Barriers to implementation

One of the primary barriers to implementing a within-network sharing policy, is generating buy-in and motivating grantees to share their outputs on the Hub by the time of publication. While some grantees regularly share their outputs on the Hub, others wait until annual progress reports are due, a time when they know that ASAP leadership are assessing their team’s progress. In an effort to assist grantees, we have recently started reminding grantees of this requirement and provide a list of research outputs that have been identified in manuscripts and need to be added to the Hub as a gentle reminder to grantees about our policy (Figure 2).

**Figure 2.**
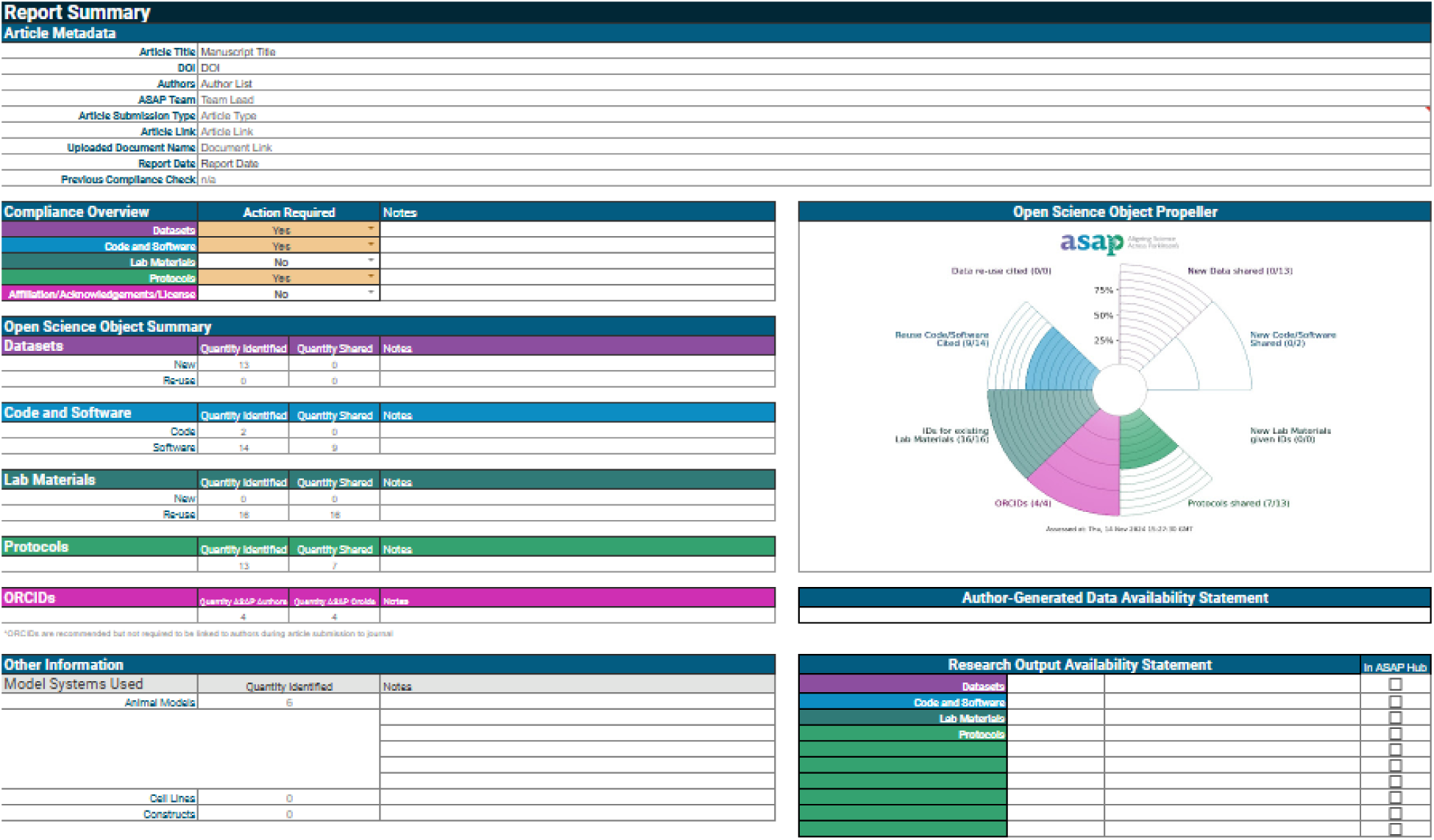
Example summary page of a deidentified ASAP compliance report. DataSeer generates compliance reports in Google sheets that provide an itemized list of all research outputs and inputs, if and how they are shared, and what still needs to be shared. The compliance report also provides information to grantees about licensing, ORCIDs and affiliations, and whether or not research outputs have been shared on the ASAP Hub.

### Costs associated with grantee network infrastructure

Creation of a virtual grantee platform could be accomplished using free, online tools like [Notion](https://www.notion.com/?gad_source=1&gclid=Cj0KCQiAx9q6BhCDARIsACwUxu6ZlCMl4hPx1KN1FxHX1P-vJsc5c7IBRj0QOI1x1KGBVmYfdynKbKMaAifTEALw_wcB) where the investment may be in the thousands for additional add on features. Funders may also choose to develop their own custom platform which could cost up to millions of dollars for development and maintenance. The use of a virtual grantee platform is not necessary in order to implement an open science policy, however, it can provide benefits to the grantee and provides an important means of tracking grant-related outputs.

## Lessons Learned

The ASAP Initiative has yielded many lessons learned with regard to the implementation of and costs associated with an open science policy so far. Importantly, we have found that policy enforcement, education and training, and utilization of ASAP-funded project managers are key to successful implementation of a progressive open science policy.

### Policy Enforcement

The [ASAP Open Science Policy](https://parkinsonsroadmap.org/open-science-policy/#) consists of five key requirements, with which every ASAP-funded manuscript must comply. To ensure compliance, the ASAP Open Science Policy requires that manuscript drafts must be sent to the ASAP Open Science Team no later than the time of posting a preprint to undergo a comprehensive review process. We partner with [DataSeer](https://dataseer.ai/) who use artificial intelligence and natural language processing to catalog research inputs and outputs into compliance reports. DataSeer generates a compliance report for every ASAP-funded manuscript, which is subsequently reviewed by the ASAP Open Science Team and shared with authors (Figure 2).

This partnership with DataSeer represents a large annual expenditure. Additionally, enforcement requires a team of scientists who can evaluate manuscripts and/or DataSeer compliance reports for openness and accuracy, which represents a potentially substantial labor cost depending on the influx of manuscripts being reviewed. At the beginning of the ASAP grant period, there were no designated 100% full-time effort (FTE) employees for the open science workflow. As of January 2024, however, the ASAP Open Science Team has grown to three staff (2 × 100% FTE, 1 25% FTE). These labor costs are critical to ensure that our open science policy is upheld.

The compliance review process is critical for enforcement of the ASAP Open Science Policy. Between 2021-2024, papers were more compliant with the ASAP Open Science Policy if they underwent a compliance review by the ASAP Open Science Team (Table 5). In 2024, publications that did not go through a compliance review were on average 33% compliant with the ASAP Open Science Policy (Table 5). By contrast, publications that had been through a previous compliance review were on average 76% compliant (Table 5). The partnership with DataSeer is a strategic investment towards ensuring enforcement of the ASAP Open Science Policy.

**Table 5.** Impact of ASAP compliance review on openness of final publications. Compliance was calculated as a percentage. Total research inputs and outputs shared across all papers divided by total research inputs and outputs identified across all papers. Note that each research input and output type was weighted the same for this calculation, for example sharing a protocol and sharing a dataset were weighted the same. Data was pulled only from ASAP-funded publications that have been through a compliance review and were quality controlled by ASAP Staff. Percentages are rounded to the nearest whole number.

| Year | Publication Version = 1 | Publication Version >1 |
| --- | --- | --- |
| 2021 | 0% | 25% |
| 2022 | 8% | 49% |
| 2023 | 14% | 61% |
| 2024 | 33% | 76% |
| All Years | 21% | 62% |

In addition to the compliance review process, funders should consider possible labor costs associated with tracking preprints and publications. ASAP becomes aware of preprints and publications through three main mechanisms: 1) a grantee has emailed the manuscript to the ASAP Open Science Team, 2) a grantee has posted the preprint or publication on the ASAP virtual Hub, or 3) we discover the preprint or publication via automated alerts from bioRxiv, medRxiv, or OA.Report. [OA.Report](https://oa.works/), which costs around $10,000 annually, is a service which identifies ASAP-funded preprints and publications through a combination of automated and manual searches of online databases (e.g., OpenAlex, bioRxiv, and medRxiv). It is the primary means by which we capture ASAP-funded publications to be added to our tracking system and a strategic investment towards enforcing our open science policy.

### Incentivization and Education

Incentivizing grantees to share research outputs and providing educational resources has been key to grantee compliance with the ASAP Open Science Policy. Grantees awarded ASAP funding agree to follow our open science requirements during the contracting phase. However, we recognize that proactively following the policy is not trivial. So, we incentivize compliance by paying APCs and through recognition of outstanding efforts towards open science via the ASAP CRN [Open Science Champion](https://parkinsonsroadmap.org/open-science-champions/#) award.

In addition, current grantees are aware that if they apply for future funding opportunities, their past record of compliance with the ASAP Open Science Policy will be a factor in the funding decision. We provide feedback to grantees on how they are doing in regards to open science compliance on an annual basis. Additionally, we provide a variety of educational resources to grantees about open science, including a suite of Open Science Guides and an itemized Policy Handbook, available on Zenodo. The ASAP Open Science Team also provides open science consultation calls and private workshops to grantees. Project managers part of ASAP grantee teams receive additional training through an hour-long onboarding process that covers the policy requirements and the compliance review workflow. We also publish [blog posts](https://parkinsonsroadmap.org/newsroom/#) that share case studies and stories behind the requirements of the ASAP Open Science Policy, in an effort to generate buy-in for these practices.

### Project Managers

The ASAP CRN requires that each research team hire a project manager as part of their awarded ASAP grant budget. Project managers are responsible for facilitating communication between labs, and disseminating information from ASAP to their team about new guidance and updates to the ASAP Open Science Policy. Project managers play a key role in ensuring compliance with the ASAP Open Science Policy, providing guidance and expertise on where and how to share data, generation of key resource tables, writing and sharing of protocols, etc. Within the compliance workflow, project managers are tasked with sharing all team manuscripts with the ASAP Open Science Team and are responsible for keeping their team’s outputs on the Hub up to date.

## Conclusion

The ASAP Initiative is committed to accelerating the pace of discovery in Parkinson’s disease research through three foundational principles: supporting collaboration, generating research enabling resources, and data sharing. These principles are built upon a strong, comprehensive open science policy. The [ASAP Open Science Policy](https://parkinsonsroadmap.org/open-science-policy/#), as described throughout this paper, has several key requirements, each of which have varying financial and labor costs. Here, we have outlined the benefits and costs associated with our policy so that other funding agencies can take these into consideration as they design their own open science policies. Overall, costs that a funder will incur while operationalizing and implementing an open science policy will vary depending on policy requirements respective to the funder. When designing and implementing an open science policy, funders should consider which aspects align with their resources, budget, and capacity. Nevertheless, open science policies, whether fully comprehensive or selectively implemented, can foster greater transparency, collaboration, and accessibility in research, contributing to the advancement of scientific knowledge and accelerating the pace of discovery.

## Acknowledgements

The authors would like to thank Cornelis Blauwendraat and Randy Schekman for their feedback and review of the manuscript. We would also like to thank Matt Lewis, Andrew Koemeter-Cox, Hetal Shah, Alexandra Vaina, and Shalini Padmanabhan on the ASAP Scientific Facilitation team, Tim Vines of DataSeer and Lenny Teytelman of protocols.io for helping us shape our open science ecosystem implementation. We also would like to thank all ASAP grantees for participating in our Open Science policies.

## Availability Statement

The data generated in this study can be found on Zenodo (https://zenodo.org/records/14262739). No code was generated for this study; all data cleaning, preprocessing, analysis, and visualization was performed using Excel or GraphPad Prism. The ASAP Open Science Policy Handbook (https://zenodo.org/records/13769766) and the ASAP Open Science guides (https://zenodo.org/records/13769802) can both be found on Zenodo.

